# The Tec kinase ITK differentially optimizes NFAT, NF-κB, and MAPK signaling during early T cell activation to regulate graded gene induction

**DOI:** 10.1101/2020.11.12.380725

**Authors:** Michael P. Gallagher, James M. Conley, Pranitha Vangala, Andrea Reboldi, Manuel Garber, Leslie J. Berg

## Abstract

The strength of peptide:MHC interactions with the T cell receptor (TCR) is correlated with the time to first cell division, the relative scale of the effector cell response, and the graded expression of activation-associated proteins like IRF4. To regulate T cell activation programming, the TCR and the TCR proximal kinase ITK simultaneously trigger many biochemically separate TCR signaling cascades. T cells lacking ITK exhibit selective impairments in effector T cell responses after activation, but under the strongest signaling conditions ITK activity is dispensable.
To gain insight into whether TCR signal strength and ITK activity tune observed graded gene expression through unequal activation of disparate signaling pathways, we examined Erk1/2 activation and NFAT, NF-κB translocation in naive OT-I CD8^+^ cell nuclei. We observed consistent digital activation of NFAT1 and Erk-MAPK, but NF-κB displayed dynamic, graded activation in response to variation in TCR signal strength and was tunable by treatment with an ITK inhibitor. Inhibitor-treated cells showed dampened induction of AP-1 factors *Fos* and *Fosb*, NF-κB response gene transcripts, and survival factor *Il2* transcripts. ATAC-seq analysis also revealed genomic regions most sensitive to ITK inhibition were enriched for NF-κB and AP-1 motifs. Specific inhibition of NF-κB during peptide stimulation tuned expression of early gene products like c-Fos. Together, these data indicate a key role for ITK in orchestrating optimal activation of separate TCR downstream pathways, specifically aiding NF-κB activation. More broadly, we revealed a mechanism by which variation in TCR signal strength can produce patterns of graded gene expression in activated T cells.

## INTRODUCTION

After T cell receptor (TCR) triggering, a single naïve CD8+ T cell has the potential to expand into millions of daughter effector cells which use cytolytic factors to eradicate virus-infected cells. The strength of the interaction between the TCR and cognate peptide:MHC molecules on antigen presenting cells (APCs) controls the rapidity the response and the ultimate scale of the effector pool. Stronger-affinity TCR interactions lead to prolonged periods of proliferation and longer times of engagement with APCs, thereby producing larger pools of CD8^+^ effector T cells (1–3).

TCR triggering and proximal signaling events exhibit noisy, switch-like behavior (4). The kinetic proofreading model of TCR signal initiation posits that ligand discrimination is guarded by the accumulation of rate-limiting signaling intermediates which elicit committed activation of transcription factors. Strong peptide:MHC ligands bind frequently with the TCR and overcome these rate-limiting steps, while weak ligands bind less frequently and are less likely to accumulate signaling intermediates (4–6). Individual downstream transcription factors pathways also display digital signaling behaviors. TCR-mediated store operated calcium (Ca^2+^) entry (SOCE) and subsequent nuclear factor of activated T cells (NFAT) activation display probabilistic, digital triggering corresponding to the dose of peptide:MHC (7–10). We recently demonstrated that NFAT1 nuclear translocation in response to TCR stimulation is a digital process, as Ovalbumin (OVA) peptide concentration modulates the frequency of responder cells within a naïve clonal OT-I T cell population, without affecting the amount of NFAT1 protein measured in individual responding cell nuclei (10). Similar bimodal behavior is described for extracellular signal-regulated kinase (Erk) activation, as OVA peptide dose carefully tunes the number of digitally activated Erk responders in a pool of naïve OT-I cells (11, 12). The behavior of these pathways is enforced through feedback mechanisms (Erk) and rapid regulation of phosphorylation states (NFAT) (8, 12). Nuclear factor (NF)-κB signaling in T cells has also been observed to behave digitally under some conditions (13). Following TCR engagement, these separate pathways activate in concert. However, careful observation of the simultaneous and relative behavior of each pathway in T cells in response to peptide stimulation is lacking (14).

Peptide concentration and avidity contribute to the probability of digital TCR triggering in individual cells, but at the same time these same variables tune the graded expression of important effector-associated factors, notably interferon regulatory factor (IRF)4 and CD25 (10, 12, 15, 16). This disconnect prompted us to question how graded gene expression patterns emerge from digital signaling events. Interleukin-2-inducible T cell Kinase (ITK) is a critical component for optimal activation of phospholipase-C-gamma (PLCγ), the enzyme that cleaves the membrane-embedded phosphatidylinositol bisphosphate (PIP_2_) into equimolar amounts of two second messengers: inositol triphosphate (IP_3_) and diacylglycerol (DAG). IP_3_ and DAG are responsible for robust activation of downstream TCR signaling pathways including NFAT, Erk, and NF-κB (17, 18). Although TCR signaling is not completely abolished in the absence of ITK, T cells from *Itk*-/- mice show inefficient intracellular Ca^2+^ flux and have notable defects in Erk phosphorylation (p-Erk) (19–22). Stimulation of naïve *Itk*^-/-^ OT-I cells or treatment with a small molecule ITK inhibitor limited the induction of IRF4 (15), indicating that ITK may not regulate gene expression in an all-or-none fashion, but rather act as a rheostat to carefully tune TCR signaling.

To determine whether ITK differentially regulates digital TCR responses to tune graded gene expression, we measured simultaneous NFAT1, NF-κB and Erk activation in single naïve OT-I T cells stimulated with peptides of variable affinity with or without use of an inhibitor of ITK (and resting lymphocyte kinase, RLK). We found that NFAT1 and Erk activation reliably responded digitally, including in weakly stimulated cells, however each pathway has a different sensitivity to ITK/RLK inhibition. Importantly, NF-κB activation occurred incrementally; cells that digitally triggered NFAT1 had graded amounts of NF-κB activation that scaled with peptide affinity and was sensitive to ITK/RLK inhibition. We also measured the immediate transcriptional response following different TCR signaling conditions which revealed expression of NF-κB target transcripts and AP-1 factors were most sensitive to ITK/RLK-inhibition. Regions of changing DNA accessibility most sensitive to ITK/RLK inhibition were also enriched for NF-κB and AP-1 binding motifs. Inhibition of NF-κB activation in stimulated OT-I cells altered TCR-induced gene expression in a pattern that mirrored the effects ITK/RLK inhibition. Together, these data underscore a role for ITK as an amplifier of TCR signaling; critical to tune NF-κB signaling in digitally-activated naïve CD8^+^ T cells.

## MATERIALS AND METHODS

### Mice

Mice were bred and housed in a specific pathogen-free facility at the University of Massachusetts Medical School (Worcester, MA) in accordance with Institutional Animal Care and Use Committee guidelines. OT-I transgenic *Rag1*-/- mice (B6.129S7-*Rag1^tm1Mom^* Tg(TcraTcrb)1100Mjb N9+N1) and C57BL/6 wild type mice were purchased from Taconic Biosciences. CD45.1+ (B6.SJL-PtprcaPep3b/BoyJ) mice were purchased from The Jackson Laboratory. Unless otherwise noted, experimental cohorts consisted of age and sex-matched littermates aged 6-12 weeks.

### Stimulation of CD8+ T cells

Freshly harvested OT-I *Rag1*-/- mouse splenocytes were pooled, RBC lysed, and enriched for CD8+ cells with an EasySep™ negatively selective magnetic isolation kit (STEMCELL Technologies). OT-I cells prepared for use in nuclei isolation experiments were then treated with CellTrace™ Violet reactive dye (Invitrogen) for 20 minutes to fluorescently label cells (including nuclei). OT-I cells were cultured at 2 × 10^5^ cells per well (unless otherwise noted) and incubated with or without 50 nM (or otherwise noted) ITK/RLK inhibitor PRN694 (Principia Biopharma) for 30 minutes 37°C. In some experiments, OT-I cells were incubated with or without IKK inhibitor IKK-16 (Sigma) or MEK inhibitor PD325901 (Tocris Bioscience). For antigen-presenting cells, RBC-lysed splenocytes harvested from wild type mice were cultured at 4 × 10^5^ per well and incubated with indicated concentrations of OVA “N4” peptide (SII**N**FEKL), altered OVA “T4” peptide (SII**T**FEKL), altered OVA “G4” peptide (SII**G**FEKL) (21^st^ Century Biochemicals) for 30-60 minutes at 37°C. OT-I cells and peptide loaded splenocytes were then combined and incubated at 37°C for specific times. For cell preparations used for molecular analyses (e.g. RNA-seq and ATAC-seq analyses), splenocytes from CD45.1+ wild type mice were used as peptide presenting cells for easy exclusion from CD45.2+ OT-I cells via cell sorting.

### Nuclei isolation

To measure translocation of nuclear proteins, we isolated cell nuclei after stimulation for fixation and subsequent flow cytometry analysis. To do this, we utilized a sucrose buffer-based protocol that we and others have previously published (7, 10). To summarize, stimulated cells were pelleted and washed with 200 μL of ice cold “Buffer A” containing 320 mM sucrose, 10 mM HEPES (Life Technologies), 8 mM MgCl_2_, 13 EDTA-free cOmplete Protease Inhibitor (Roche), and 0.1% (v/v) Triton X-100 (Sigma-Aldrich). After 15 min on ice, the plate was spun at 2000 × *g* and 4°C for 10 min. This was followed by 2X 200 μL washes with “Buffer B” (Buffer A without Triton X-100) at 2000 × *g* and 4°C.

### Antibodies and flow cytometry

Stimulated cells and isolated cell nuclei were fixed and permeabilized with the Foxp3 / Transcription Factor Staining Buffer Set (eBioscience), except cells used for anti-p-Erk1/2 analysis, which were fixed with 4% paraformaldehyde (Electron Microscopy Services) and permeabilized with 90% ice-cold methanol (FisherSci). Fluorescently-labeled flow cytometry antibodies against IRF4 (3E4), CD69 (H1.2F3), and Egr2 (erongr2) were purchased from eBioscience. Antibodies against CD8a (53-6.7), CD8b (53-5.8), CD25 (3C7), CD90.2 (53-2.1), p-Erk1/2 (4B11B69) were purchased from BioLegend. Antibodies against NFAT1 (D43B1), NF-κB p65 (D14E12), c-Fos (9F6), and c-Myc (D84C12) were purchased from Cell Signaling Technology. Anti-CD45.1 (A20) was purchased from BD Pharmingen. PE-conjugated F(ab’)2-goat anti-rabbit IgG (H+L) cross-adsorbed secondary antibody was purchased from Invitrogen.

### Cell Sorting

Samples of stimulated CD45.2+ OT-I cells and CD45.1+ wild type splenocytes mixtures were stained with CD8a and CD45.2 antibodies and 7-AAD and OT-I cells were sorted (BD FACSAria) into 100% FBS and pelleted.

### RNA-seq library preparation

Total RNA from ~300,000 OT-I cells per sample was collected with the RNeasy micro kit (Qiagen) with a 15-minute on-column DNase digestion (Qiagen) to remove genomic DNA. Total RNA quality and quantity was determined with fragment analysis (University of Massachusetts Medical School Molecular Biology Core Lab) and Qubit Fluorometer (Invitrogen) analysis. cDNA libraries were generated following a modified paired-end SMART-seq protocol (23, 24). Briefly, at least 20 ng of input RNA was used for reverse transcription with SMARTscribe reverse transcriptase (Clontech). Whole transcriptome amplification (WTA) was performed with Advantage 2 polymerase (Takara Bio). WTA reactions were monitored with qPCR to determine optimal cycle number. WTA libraries were then sized-selected with AMPure XP DNA SPRI beads (Beckman Coulter), tagmented with Tn5 transposases (Illumina Nextera XT), barcoded, and amplified with cycle number determined via qPCR monitoring. Final libraries were further size-selected with SPRI beads to an average size of 300-500 bp and quality was assessed with fragment analysis and Qubit analysis. Libraries were pooled and sequenced on a NextSeq 500 sequencer (Illumina).

### Processing and analysis of RNA-seq reads

Adapter sequences were trimmed from quality raw sequencing reads with Trimmomatic-0.38 (25) and then aligned to mouse ribosomal RNA with Bowtie2 v2.3.2 (26). Unaligned reads were retained, and gene expression was estimated (transcripts per million and expected counts) with RSEM v1.2.29 (27) configured to align to a mm10 RefSeq transcriptome with Bowtie2 v2.3.2. Samples were filtered to retain expressed genes (expected counts > 200) and batch effects between replicates were corrected with limma v3.42.2 (28). Differential expression analysis was performed with DESeq2 v1.26.0 (29) to identify induced genes (stimulated conditions vs. unstimulated controls) or condition-specific changes in expression (untreated vs. PRN694 or N4 vs T4 OVA peptides). Hierarchical clustering and k-means clustering of differentially expressed genes was performed within R v3.5. Heatmap visualizations of gene clusters were drawn with ComplexHeatmap (30).

### Gene ontology

We utilized the R package msigdbr v7.0.1 to compare clusters of differentially expressed genes with the Molecular Signatures Database (MSigDB) Hallmark (H) and Immunologic (C7) gene sets (31–33). The top five terms with FDR <0.05 were displayed in the Results.

### ATAC-seq library preparation

Precisely 50,000 stimulated OT-I T cells were FACS sorted at the same time as RNA-seq samples and pelleted in 100% FBS. ATAC-seq libraries were generated similarly as described in Buenrostro et al., 2013 (34). Briefly, cell nuclei were isolated and transposed with 8 μL Tn5 (Illumina Nextera) at 37°C for 60 minutes. DNA fragments were isolated with a Clean and Concentrator Kit (Zymo) and then Illumina barcoded and amplified with NEBNext High-Fidelity 2x PCR Master Mix (New England Biolabs). A portion of the reaction was performed as a separate qPCR reaction to determine ideal cycle number. Samples were then size selected with SPRI beads to include fragments up to 450 bp, ensuring to maintain small (<200 bp), nucleosome-free fragments. Quality of final ATAC-seq libraries was assessed with fragment analysis and Qubit analysis. Libraries were then pooled and sequenced on an Illumina NextSeq 500.

### Alignment and processing of ATAC-seq reads

Adapter sequences were trimmed from raw sequencing reads with Cutadapt v1.3 (35) and then aligned to the mouse genome (mm10) with Bowtie2 v2.1.0 with the parameter -X 2000. PCR duplicates were removed with Picard’s markDuplicates v2.17.8 and aligned reads were sorted and filtered with SAMtools v1.4.1 (36). For visualization of fragment coverage TDFs were generated with IGVTools v2.3.31 (37, 38).

### Peak calling

Aligned ATAC-seq reads were trimmed to 29 bases closest to the Tn5 cut site with bedtools v2.26.0 (39) and then peaks were called with MACS2 v2.1.1 (40) using parameters --bw 29 --nomodel -q 0.0001. Summits of called peaks across all samples were merged and “slopped” ±100bp to create a master peak reference file using bedtools v2.26.0. Peaks were annotated with names of closest genes using a mouse OT-I TSS BED file and bedtools v2.26.0 (closest -D ref -t all). Peaks with summits within 500 bp from TSSs were labeled “promoter” peaks and summits further than 500bp were labeled “enhancer” peaks. Peak coverage for each ATAC-seq sample was calculated using bedtools v2.26.0 (intersect; coverage).

### Differential Peak Analysis

Calculated peak coverage values for two ATAC-seq replicate experiments were used as input for differential analysis using DESeq2 v1.26.0. All differential peaks (compared to unstimulated controls) were clustered using either hierarchical or k-means clustering methods within R. Heatmaps were generated with ComplexHeatmap.

### Motif enrichment

Each cluster of annotated differentially accessible peak regions was tested for de novo motif enrichment using HOMER v4.10.3 (findMotifsGenome.pl -size 200) with background comprised of peak regions found in all other clusters (41).

### Genomic Visualizations

Genomic tracks were created by plotting BigWig files with help of Gviz (42) PMA and ionomycin ATAC-seq tracks and NF-κB ChIP-seq tracks were extracted from publicly available sources (43, 44).

## RESULTS

Signaling pathway activation after TCR engagement has largely been described as digital, where TCR triggering is guarded by a threshold for activation and downstream responses of NFAT, NF-κB and MAPK then “switch on” together (9, 12, 13). The probability that a TCR stimulus will digitally activate an individual naïve T cell within a stimulated clonal population is measured by examining the fraction of cells that upregulate the cell surface marker CD69 (11, 12, 16, 18). Cells experiencing weakened TCR stimulation may sufficiently switch on digital CD69 expression but fail to maximally upregulate important effector-associated factors like IRF4 (16). Additionally, CD69 expression does not explicitly guarantee that a T cell will be sufficiently stimulated to commit to clonal expansion and effector programming (45, 46). Thus, there are signaling behaviors underlying digital triggering that generate divergent gene expression programs and ultimately contribute to a naïve T cell’s fate. We hypothesized that the activity of the TCR proximal tyrosine kinase ITK is sensitive to variable amount of TCR stimulation in order to tune the intensity of downstream signaling pathways and the graded transcription of a subset of genes induced in digitally activated cells.

### ITK/RLK inhibition differentially dampens gene expression in activating OT-I cells

To test whether tunable ITK activity modulates TCR signaling during activation, we measured gene expression in CD8^+^ T cells treated with a covalent small molecule inhibitor (PRN694) (47). PRN694 is a compound highly selective for the active site of both ITK and Resting Lymphocyte Kinase (RLK); RLK is a kinase co-expressed in naïve T cells structurally similar to ITK without clear function in TCR signaling (48). Thymically-derived *Itk^-/-^* cells do not develop normally (22). Thus, PRN694 allows for titrated control of ITK/RLK activity in naive wild-type OT-I cells and assures both untreated and treated naïve cell transcriptional states are similar before stimulation. To regulate TCR signaling via differential ligand stimulation, we utilized OVA peptide plus altered peptide ligands, where graded peptide potency is achieved using residue substitutions within the native ‘SII**N**FEKL’ OVA peptide (N4) (2). After stimulation with weaker affinity ‘SII**T**FEKL’ (T4) altered-OVA peptide, a similar number of ITK/RLK inhibitor-treated and untreated OT-I CD8^+^ T cells upregulated CD69, but inhibitor-treated cells exhibited dampened expression of IRF4 (Fig. 1A). This confirmed that the ITK/RLK inhibitor has a differential effect on specific activation-induced genes, similar to studies examining *Itk-/-* OT-I cells (16). Inhibitor-treated T4-stimulated cells also displayed less proliferative potential measured at 48hrs, with a large percentage of cells remaining undivided (Fig. 2B). Thus, under these conditions ITK/RLK activity is dispensable for switch-like induction of the activation marker CD69, but critical in tuning the intensity of other important genes that govern T cell activation programming and division.

**Figure 1.**
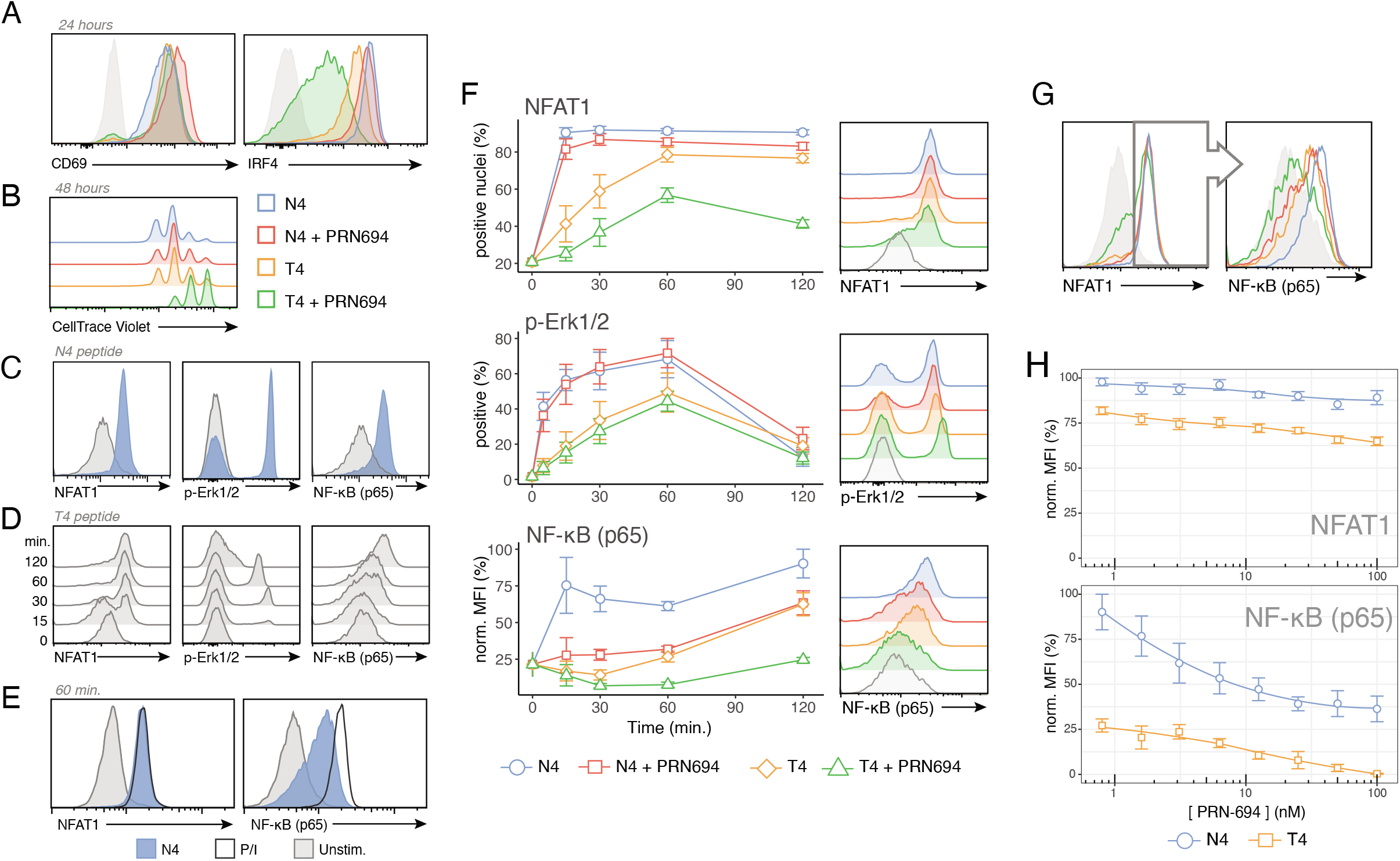
TCR signal strength and PRN694 regulate graded NF-κB activation within digitally NFAT1 and p-Erk active cells. (**A-B**) Histograms depicting CD69, and IRF4 expression after 24 hours (A) or CellTrace Violet fluorescence after 48 hours (B) in OT-I cells stimulated with APCs plus 100 nM of indicated peptide, with or without 50 nM ITK/RLK inhibitor PRN694. (**C**) Histograms of NFAT1 and NF-κB fluorescence in isolated OT-I nuclei or p-Erk1/2 fluorescence in OT-I cells after stimulation with APCs plus 100 nM N4 peptide stimulation for 1 hour. (**D**) Histograms of NFAT1 and NF-κB fluorescence in isolated OT-I nuclei or p-Erk1/2 fluorescence in OT-I cells after stimulation with APCs plus 100 nM T4 peptide for the indicated times. (**E**) Histograms of NFAT1 and NF-κB (p65) fluorescence within OT-I nuclei isolated after 1-hour stimulation with APCs plus 100 nM N4 OVA peptide or PMA (10 ng/mL)/Ionomycin (1 μM) addition. (**F**) Line plots depicting either NFAT1 or NF-κB fluorescence within OT-I nuclei or p-Erk fluorescence in OT-I cells over 2-hour stimulation with APCs plus 100 nM of indicated peptide with or without 50 nM PRN694. Histograms to the right represent nuclear or whole cell fluorescence patterns after 60 minutes of stimulation. (**G**) Histograms depicting NF-κB (p65) fluorescence within NFAT1+ nuclei after 1-hour stimulation in conditions as in A, B and E. (**H**) Line plots of change in normalized NFAT1 MFI (%) or normalized NF-κB (p65) MFI (%) within NFAT1+ OT-I nuclei after 1-hour stimulation with APCs plus 100 nM N4 or T4 OVA peptide, in the presence of titrated concentrations of PRN694. Histograms shown are representative of 3 or more independent experiments.

**Figure 2.**
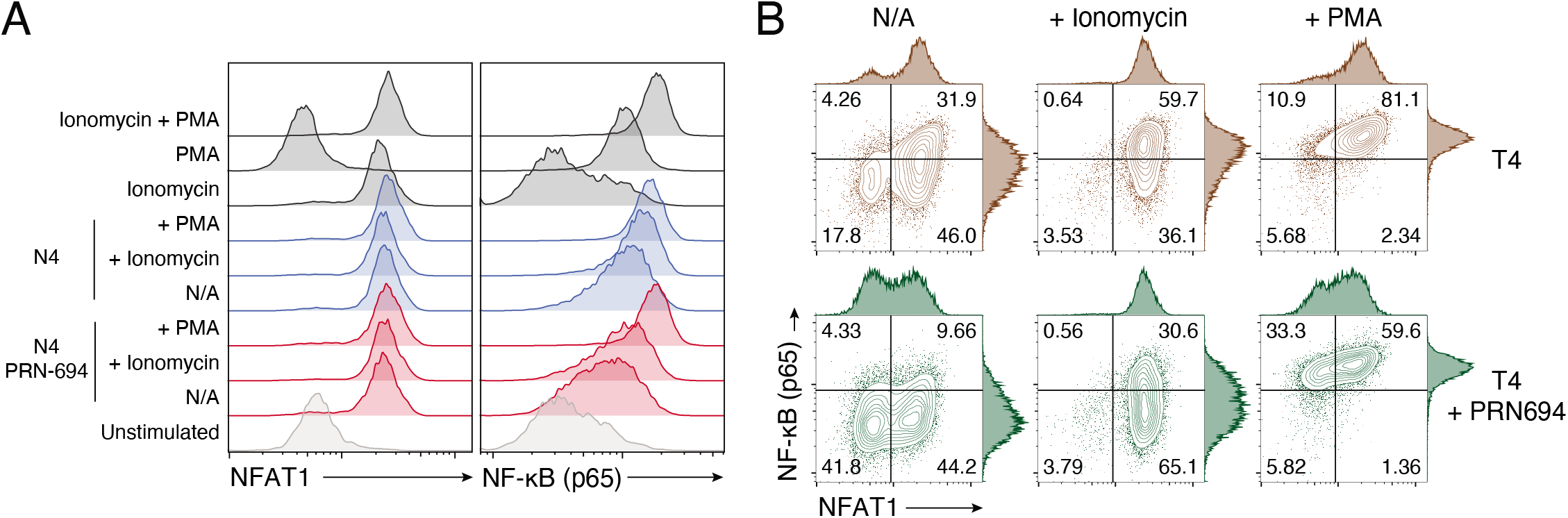
PMA and ionomycin supplementation during peptide stimulation reveals synergistic NF-κB signaling activation. (A) Histograms depicting NFAT1 and NF-κB (p65) activation during 1 hour 100 nM N4 stimulation with (red) or without (blue) 50 nM of PRN694 or received no peptide stimulation (dark grey). 1 μg/mL ionomycin or 12.5 ng/mL PMA was supplemented to wells were indicated. (B) FACS contour plots with adjust histograms comparing NF-κB (p65) and NFAT1 fluorescence in after the same stimulation conditions as in A, but with 100 nM T4 peptide stimulation.

### NF-κB activation is tunable in digitally activated naive OT-I cells

Pathways downstream of the TCR critical for robust T cell activation are highly dependent on the activation of NFAT, NF-κB, and AP-1 transcription factor families (18, 49, 50). We hypothesized that each pathway may have different sensitivity to varied amounts of upstream TCR stimulation or ITK activity and differential signaling patterns could contribute to graded expression of activation associated factors. Naïve cells have NFAT and NF-κB factors sequestered in the cytoplasm such that TCR engagement induces their rapid translocation to the nucleus (50, 51). To measure activation of NFAT and NF-κB in single cells, we performed flow cytometry with stimulated OT-I nuclei, as described previously (10). This technique allowed us to quantify both the proportion of OT-I cells responding to stimulation as well as the relative abundance of each factor within individual stimulated nuclei.

After strong peptide:MHC stimulation, we observed that NFAT1 and NF-κB (p65) quickly translocated to the nucleus (Fig. 1C). Erk-MAPK signaling was also rapidly activated in nearly all stimulated OT-I cells, as measured by conventional phospho-Erk1/2 (p-Erk) fluorescence. These rapid responses confirmed previously reported “switch-like” signaling behavior after TCR engagement (7, 10–13). However, in response to weaker peptide stimulation, differences between pathways became more evident. Stimulation with T4 altered OVA peptide produced slower accumulation of the OT-I population exhibiting nuclear NFAT1 and p-Erk, and also reduced the maximum proportion of NFAT1 positive nuclei by 2 hours of stimulation from ~90% to 75% (Fig. 1D-F). Importantly, we observed minimal differences in NFAT1 or p-Erk MFI within responding nuclei or cells, indicating that these pathways remained digitally triggered even under weaker signaling conditions (Fig. 1D). In contrast, TCR stimulation with T4 peptide failed to optimally activate NF-κB, as evident by both the reduced number of responder cells and, by a lower intensity of NF-κB fluorescence in single OT-I nuclei (Fig. 1D). Intermediate intensity of nuclear NF-κB-p65 suggested that TCR control of the NF-κB pathway did not behave digitally, as was observed for NFAT1 and p-Erk1/2 activation. Furthermore, compared to chemical activation with PMA and ionomycin, peptide:MHC stimulated OT-I cells activated NF-κB with suboptimal intensity, while NFAT1 activation in the same cell nuclei was similar between the two stimulation conditions (Fig. 1E). These findings demonstrated that under more physiological peptide stimulation conditions, NF-κB is more tunable to the level of TCR engagement than NFAT1 and p-Erk and revealed a dynamic range of activation states within a population of stimulated T cells.

### ITK/RLK inhibition selectively dampens NF-κB intensity in activating OT-I cells

To test whether NF-κB activation was tunable by relative levels of ITK activity, we treated OT-I cells with the ITK/RLK inhibitor PRN694 prior to stimulation with peptide:MHC, and then measured patterns of NFAT1 and NF-κB (p65) translocation or p-Erk1/2 induction. During N4 OVA peptide stimulation, ITK/RLK inhibitor treatment had little effect on NFAT1 and p-Erk1/2 activation, but led to a marked reduction in the intensity of NF-κB activation (Fig. 1F,G). Notably, NF-κB MFI was reduced in cells that had a similar amount of NFAT1 fluorescence as those from the untreated samples (Fig. 1G). NF-κB MFI was also more sensitive to incremental doses of PRN694 within NFAT1-positive nuclei (Fig. 1H). During stimulation with T4 peptide, inhibitor-treated cells had weakened NFAT1, p-Erk1/2, and NF-κB activation (Fig. 1F-H). Various concentrations of strong N4, weak T4, or even weaker G4 peptide indicated that NF-κB activation was consistently sensitive to PRN964 treatment, while NFAT1 activation was less sensitive to PRN694, especially during strong N4 stimulation conditions (Supplemental Fig. 1). We interpreted the global effect of the inhibitor during weaker signaling as dilating the temporal window during which cells “switched on” along with lowering the absolute probability, or threshold, of activation. ITK/RLK inhibition mirrored the effect of varying TCR stimulation with weakened affinity peptides.

While NFAT activation is directly influenced solely by activation of calcineurin, NF-κB can be regulated by both DAG activation of IKK complex proteins as well as Ca^2+^ activation of CaM kinases (49, 52, 53). As ITK activity is upstream of both DAG and Ca^2+^, it could possibly have multifactorial control over NF-κB. To separate DAG and Ca^2+^ components of NF-κB activation in OT-I cells, PMA or ionomycin was supplemented to wells of OT-I cultures during N4 stimulation with or without PRN694. Ionomycin addition to cultures during N4 stimulation did not modify the NFAT1^+^ fraction (which was already >90% responders) or modulate the NFAT1 fluorescence intensity, but did increase the proportion NFAT1 responders during T4 stimulation as expected (Fig. 2A-B). Ionomycin alone (in the absence of peptide stimulation) was sufficient to maximally translocate NFAT1, but did not induce appreciable NF-κB translocation (Fig. 2A). Ionomycin supplementation during N4 or T4 peptide stimulation increased NF-κB (p65) fluorescence intensity within individual nuclei, indicating the threshold of Ca^2+^ flux sufficient to translocate NFAT1 is not the same as the maximum contribution to NF-κB translocation. PMA supplementation maximized NF-κB (p65) translocation in N4 or T4 stimulated cells and easily recovered the defect in NF-κB signaling due to ITK inhibition with PRN694. These experiments demonstrated that under physiological peptide stimulation, separate downstream pathways are differently sensitive to the secondary messengers generated from the results of ITK activity. They also highlighted the synergistic nature of Ca^2+^ and DAG signaling in NF-κB activation, but showed a dominant role for amounts of DAG, rather than Ca^2+^, to tune translocation.

These data demonstrated the complex behavior of concomitant signaling pathways in response to variable TCR inputs. We concluded that pathways such as NFAT1 and MAPK require a lower threshold of TCR stimulation to activate digitally, whereas NF-κB can be triggered rapidly (and appear digitally switched) under supra-physiological TCR engagement, but normal peptide stimulation taps into a dynamic range of NF-κB activation states. Also, ITK activity is crucial in ensuring optimal activation of graded NF-κB, which sheds light on the role ITK as an amplifier of TCR signals.

### ITK/RLK inhibition dampens NF-κB-associated gene expression immediately following TCR engagement

As we observed that NF-κB activation was more sensitive to ITK/RLK inhibition, we hypothesized that variable NF-κB intensity within activated cells might contribute to immediate transcriptional control of graded gene induction. Prior to upregulation of effector genes like IL-2 and IRF4, activating CD8+ T cells undergo waves of primary and transient gene transcription (54–56). To connect observed differential signaling behavior in naïve T cells to immediate transcriptional effects, we treated OT-I cells with or without ITK/RLK inhibitor and briefly stimulated with OVA N4 and altered OVA T4 peptide presented on wild type splenocytes. We then sorted OT-I cells and measured transcript abundance with RNA sequencing.

To determine whether varying peptide affinity or modulating ITK activity regulated disparate transcriptional programming, we compared transcripts upregulated in N4 or T4 stimulated OT-I cells, with or without treatment with ITK/ RLK inhibitor at 30, 60, and 120 minutes of activation. We observed that each condition, testing different qualities of TCR signaling, induced sets of transcripts largely similar in composition, but significantly different in abundance (Fig. 3A-B, Supplemental Fig. 2, Supplemental Fig. 3). While a subset of transcripts that were identified in N4 stimulated cells were not significantly upregulated in T4 stimulated cells, we found very few genes upregulated in T4-stimulated cells that were not seen in N4 stimulated cells (Supplemental Fig. 3). We clustered genes induced over two hours into six groups (Fig. 3A-B). Gene clusters that exhibited peak expression 30 minutes after TCR contact with peptide:MHC were enriched for TCR signaling-related ontology terms, including “AP-1 signaling,” “NF-κB signaling,” and “NFAT signaling” (Fig. 3C). This suggested that TCR downstream signaling pathways directly regulate transcription of these immediate early genes. Clusters of delayed genes first detectable at times greater than 30 minutes were enriched for terms linked to T cell effector functions, “Myc targets,” and cell cycle regulation, representing secondary gene transcription responses beyond direct TCR control, and likely regulated by a mix of continuing TCR signaling and first wave gene transcription. These experiments allowed us to interrogate the immediate transcriptional response to TCR engagement and revealed that modulation of upstream TCR signal strength activates overall similar transcriptional programming.

**Figure 3.**
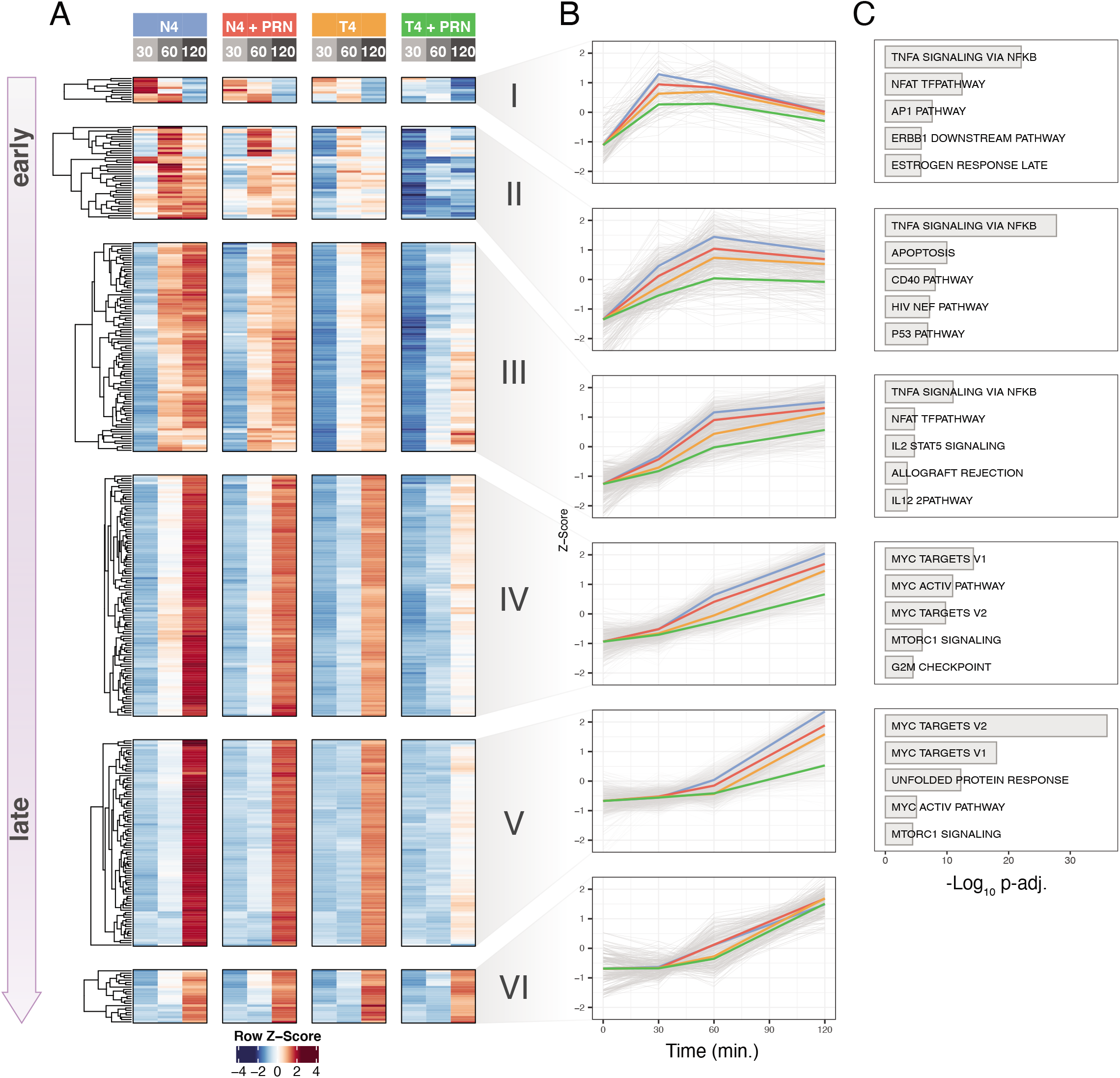
Inhibition of ITK dampens immediate TCR signaling-induced transcripts. (**A**) Heatmap depicting mean variance-stabilized transformed (VST) normalized expression values of top 357 genes (Log2 fold-change > 2, mean expression > 1000, p-adj < 0.1) induced in OT-I CD8+ T cells at three timepoints (30, 60, and 120 minutes) after stimulation with APCs plus 100 nM N4 OVA or T4 altered OVA peptide with or without 50 nM PRN694. Genes were grouped with k-means clustering into 6 clusters. (**B**) Line plots depicting Z-score of 2-hour expression time course of gene clusters identified in A. Scores for mean expression values for each cluster are grouped by conditions in A; N4 (blue), N4 + PRN694 (red), T4 (orange), T4 + PRN964 (green). Total expression data for all conditions from three replicates are drawn in gray. (**C**) Enriched MSigDB signatures in gene clusters identified in A. Log10 transformed adjusted p-values (FDR) of (up to) the top 5 terms (p-adj. ≤0.05) from both “Hallmark Gene Sets” and “Immunologic Signatures” MSigDB collections. Cluster VI did not contain gene set enrichments with these constraints. Data represent three separate biological replicate experiments each utilizing pooled splenocytes from three or more OT-I mice.

To determine whether weakened NF-κB activation during ITK/RLK inhibition differentially regulated the abundance of immediately induced transcripts, we compared differentially expressed transcripts in OT-I cells treated with or without inhibitor after 30 minutes of strong N4 peptide stimulation. Among transcripts significantly diminished by the ITK/RLK inhibitor were induced AP-1 family members (*Fos, Fosl1, Fosb, Jun*) and NF-κB response genes (*Nfkbia, Nfkbid, Nfkbiz, Tnfaip3*) (Fig. 4A). We interpret these results as reflecting the effect of weakened NF-κB signaling in ITK inhibited cells. Genes that were upregulated compared to unstimulated cells but were less sensitive to ITK/RLK inhibition included Egr family transcripts (*Egr1, Egr2, Egr3*) and the activation marker CD69 (Fig. 4A). After 2 hours of stimulation, ITK/RLK inhibitor-treated OT-I T cells also had significantly lower expression of cytokines (*IL2, Ifng*) and effector associated transcripts (*Irf4*), which is expected based on the known importance of ITK signaling in conferring robust effector functions (Supplemental Fig. 4) (57–59). Independent of TCR stimulus, most immediately-induced transcripts (*e.g. Fos, Fosb, Fosl1, Egr1, Egr2*) peaked in abundance 30 minutes after contact with peptide:MHC (Fig. 3A-B, 4A, Supplemental Fig. 4). This window of transcription is a known pattern of immediate-early gene regulation (55, 56). Our RNA-seq experiments did not suggest that ITK/RLK inhibitor caused delayed induction of these genes in strongly-stimulated OT-I cells, but effects of this type might be revealed by further single cell expression analysis approaches.

**Figure 4.**
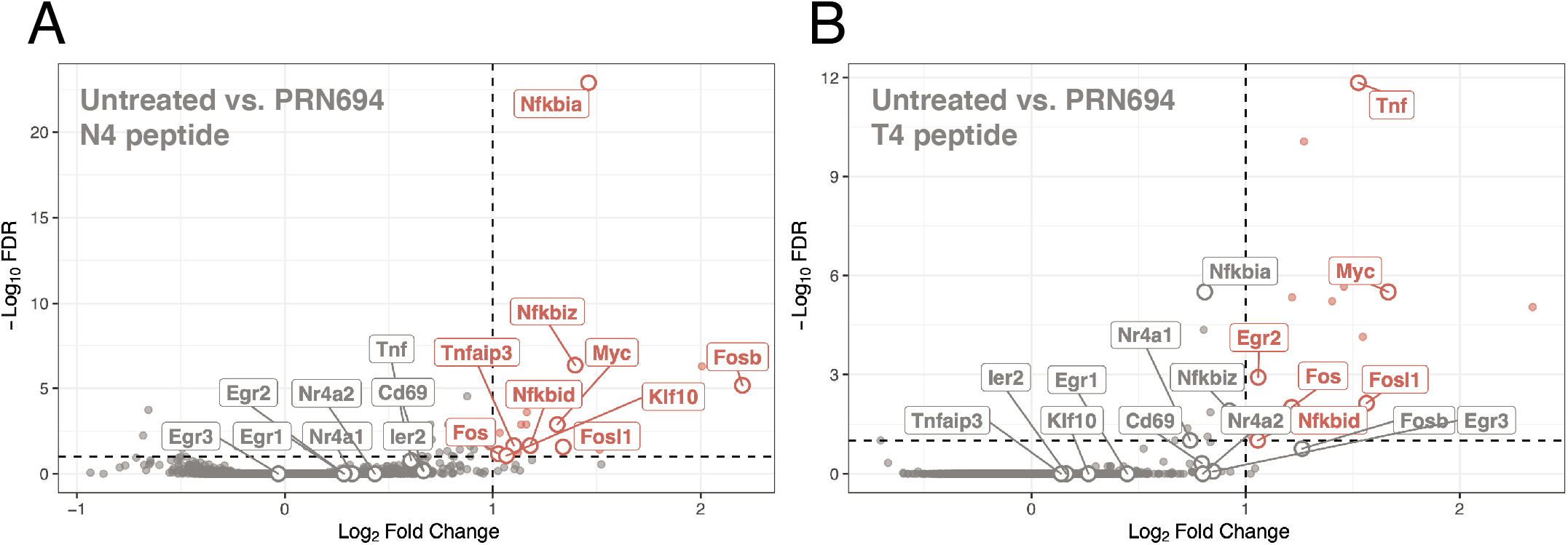
Immediate transcripts are differentially sensitive to PRN694 treatment. (**A-B**) Volcano plots identifying transcripts sensitive to 50 nM PRN694 treatment measured after 30 stimulation of OT-I cells with APCs plus 100 nM N4 peptide (A) or 100 nM T4 peptide (B). Labeled gene names represent select early-induced transcripts significantly expressed compared to unstimulated controls, extracted from either clusters I or II from Fig. 4.2. Plots depict the Log_2_ fold change of normalized expression of untreated samples over PRN694-treated samples against −Log_10_ transformed FDR values. Data points and gene names significantly above FC cutoffs (Log_2_ fold change > 1, −Log_10_ FDR <0.1) are drawn in red; genes that were not significantly differentially expressed due to PRN694 treatment are drawn in gray.

### Changes in DNA accessibility sensitive to ITK/RLK inhibitor are enriched for NF-κB and AP-1 motifs

To connect individual TCR signaling pathway behavior with ITK control of immediate gene expression patterns, we measured immediate, genome-wide DNA accessibility changes with ATAC-seq and analyzed enrichment of transcription factor binding motifs using HOMER (34, 41). Cells for ATAC-seq analysis were sorted at the same time as those used in RNA-seq experiments, to better compare accessibility changes with gene expression. ATAC-seq replicates clustered together when plotted with PCA, indicating that variance was attributed to differences in stimulation. Similar to patterns found in transcriptome analysis, the strength of the TCR stimulation or ITK inhibition did not regulate an independent set of genomic regions, but rather regulated the intensity of a shared set of activation-associated pileups (data not shown). Compared to naive, unstimulated OT-I cells, strong N4 stimulation induced the most significant differences in DNA accessibility, most evident after 120 minutes (Fig. 5A). After only 30 minutes of N4 stimulation, about 10,000 DNA regions displayed differential accessibility compared to unstimulated cells (Fig. 5B). To evaluate differential accessibility due to ITK inhibition, N4 stimulated cells with and without ITK inhibition were compared. After 30 minutes of stimulation, the most significant changes in accessibility due to PRN694 were found near many gene loci also identified in early transcriptional data including AP-1 factors (*Fosb, Jun*), NF-κB response genes (*Nfkbia, Nfkb1*), and genes encoding transcription factors important in regulating effector function (*Rel, Nfatc1, Irf4*) (Fig. 5C). This indicated that while many genomic regions change in accessibility similarly during activation, the few that were most sensitive to PRN694 also were near early genes that echoed the most sensitive gene sets found in the transcriptional analysis.

**Figure 5.**
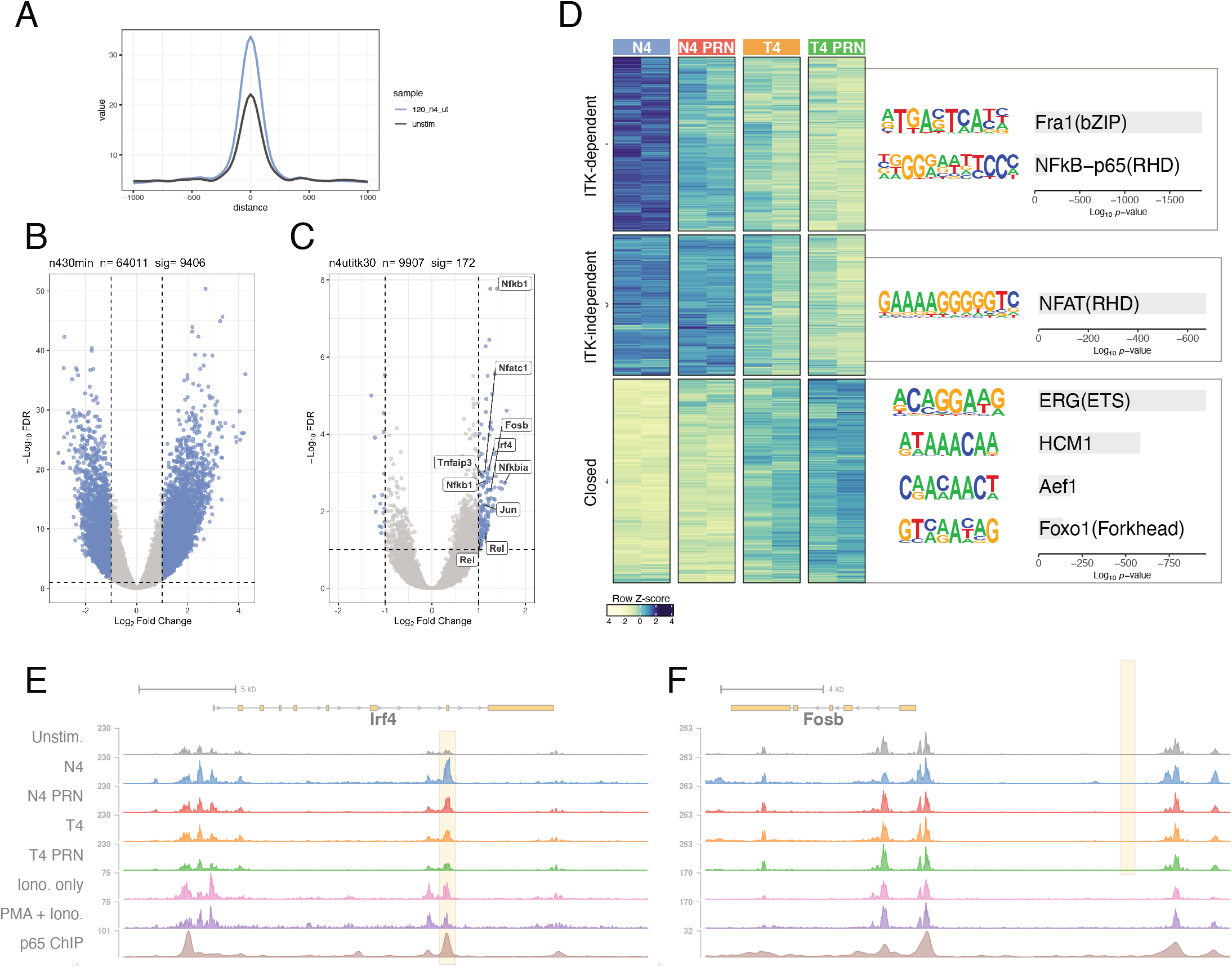
Early ITK-dependent changes in DNA accessibility are enriched for NF-κB and AP-1 binding motifs. (**A**) Distribution of ATAC-seq fragment distance from peak centers. Comparison of 120-minute stimulation with N4 peptide (blue) and unstimulated OT-I nuclei (grey). (**B**) Volcano plot of differentially-accessible genomic peak regions after 30 minutes of stimulation with N4 peptide. (**C**) Volcano plot of differentially accessible peak regions due to PRN694 treatment after 30 minutes of N4 peptide stimulation. Annotations of closest gene TSS are labeled for select regions. For B and C, significantly differentially accessible regions (Log_2_ fold change < −1 or > 1 and *p-adj* < 0.1) are drawn blue. (**D**) Heatmap and HOMER *de novo* motif enrichment analysis of *k*-means clustered Log2 transformed and row-scaled peak regions after 30 min of either 100 nM N4 or T4 peptide stimulation with or without 50 nM PRN694 (N4 untreated, blue; N4 + PRN694, red; T4 untreated, orange; T4 + PRN694, green). Displaying top (*p*-adj < 10^-50^) motifs for each cluster. (**E-F**) ATACs-seq genomic tracks for regions around *Irf4* and *Fosb* loci comparing different peptide and PRN694 stimulation conditions after 30 minutes. Also plotted are ionomycin or ionomycin + PMA stimulation ATAC-seq peaks and NF-κB ChIP-seq peaks (Mognol et al, 2017; Oh et al., 2017). Differential experimental ATAC-seq peaks are highlighted in yellow.

Similar to the RNA-seq experiments, the strength of the TCR stimulation or ITK/RLK inhibition did not regulate an independent set of genomic regions, but rather regulated the intensity of a shared set of activation associated pileups (Fig 5E-F). *k*-means clustering of all 30 minute activation-induced DNA accessibility changes (differentially-accessible compared to unstimulated control) for all sample conditions revealed regulation of regions that had increased dependency on ITK activity (Fig. 5D). Regions that were less accessible after treatment with the ITK/ RLK inhibitor were significantly enriched for AP-1 (“Fra1”) motifs and NF-κB motifs. Regions that were less sensitive to PRN694 treatment were significantly enriched for NFAT family motifs. The ATAC-seq results indicate that specific DNA regions were differentially more reliant on ITK activity for optimal regulation. Based on the motif enrichment data, we interpreted these accessibility changes to be due to, at least in part, decreased NF-κB signaling in ITK/RLK-inhibited cells.

### TCR stimulus and ITK regulate graded selective expression of early gene products

To determine whether ITK/RLK inhibitor-specific effects on immediate transcription also led to reduced protein product accumulation during activation, we measured Egr2, c-Fos, c-Myc intracellular content in stimulated cells via flow cytometry (Fig. 6A-C). Both c-Fos and Egr2 proteins were detectable within 30 minutes after OVA stimulation. Treatment with the ITK/RLK inhibitor dampened the amount of c-Fos expression in OVA-stimulated OT-I cells, but had little effect on modulating Egr2 expression. After approaching a peak expression at 60 minutes, the expression of both proteins was stable out to 6 hours. This indicated that ITK/ RLK-inhibited OT-I cells did not “catch up” in c-Fos expression during the time course of this experiment and the RNA-seq experiments measured dampened expression and not an average of asynchronous cells. Cells that expressed both CD69 and Egr2 also displayed a graded amount of c-Fos expression dependent on the TCR stimulation strength and ITK/RLK-inhibition (Fig. 6D). These results also confirmed that ITK/RLK signaling contributes to graded gene expression in response to variations in TCR signal strength.

**Figure 6.**
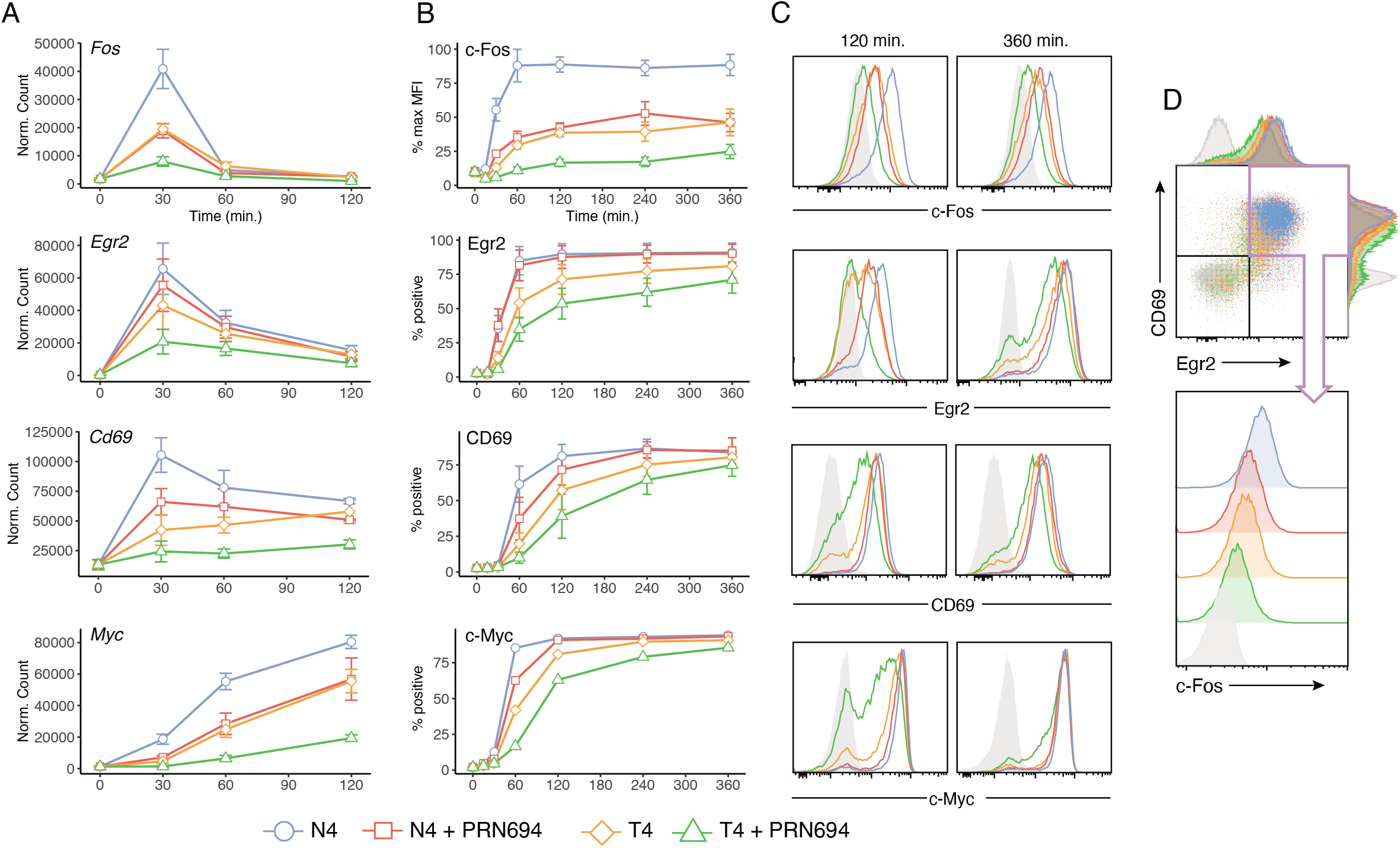
TCR signaling and ITK control graded accumulation of early gene product c-Fos. **(A)** Line plots of mean normalized expression counts of select induced transcripts measured at 0, 30, 60, and 120 min similar to Fig. 4.4. **(B-C)** Line plots (B) and representative histograms (C) of flow cytometry results for gene products of transcripts profiled in A. Experiments were conducted under similar stimulation conditions and previous figures and stimulated up to 6 hours. Results are plotted as % positive or normalized to maximum MFI (% max MFI) where indicated. **(D)** Representative histograms depicting graded c-Fos fluorescence in CD69^+^ Egr2^+^ OT I cells in response to different stimulation conditions measured after 6 hr. Labeling is consistent: N4 - blue (circles), N4 + PRN - red (squares), T4 – orange (diamonds), T4 +PRN694 – green (triangles). Data represent three biological replicates. Error bars indicate s.e.m.

### NF-κB inhibition produces graded immediate gene expression

To test whether specific inhibition of the NF-κB pathway would also produce graded induction of early genes, we treated OT-I cells with either an inhibitor of the IκB kinase (IKK) complex, IKK-16, or for comparison, a MEK inhibitor (PD325901) to control activation of MAPK (Erk1/2). As expected, neither IKK-16 nor PD325901 had an effect on NFAT1 translocation in OT-I cells (Fig. 7A-B), but treatment with IKK-16 tuned NF-κB (p65) translocation and treatment of PD325901 effectively inhibited p-Erk (Fig. 7A-B). OT-I cells treated with moderate concentration of IKK-16 or PD325901 exhibited dampened c-Fos expression (Fig. 7C-D). As expected, Egr2 expression was also dampened by PD325901 treatment, but less affected by IKK-16 treatment. At some concentrations of IKK-16, Egr2 expression is largely unchanged but c-Fos expression is reduced two-fold (Fig. 7C, E). These experiments indicate that induction of select immediate genes, such c-Fos, are more dependent on the strength of NF-κB induction during activation.

**Figure 7.**
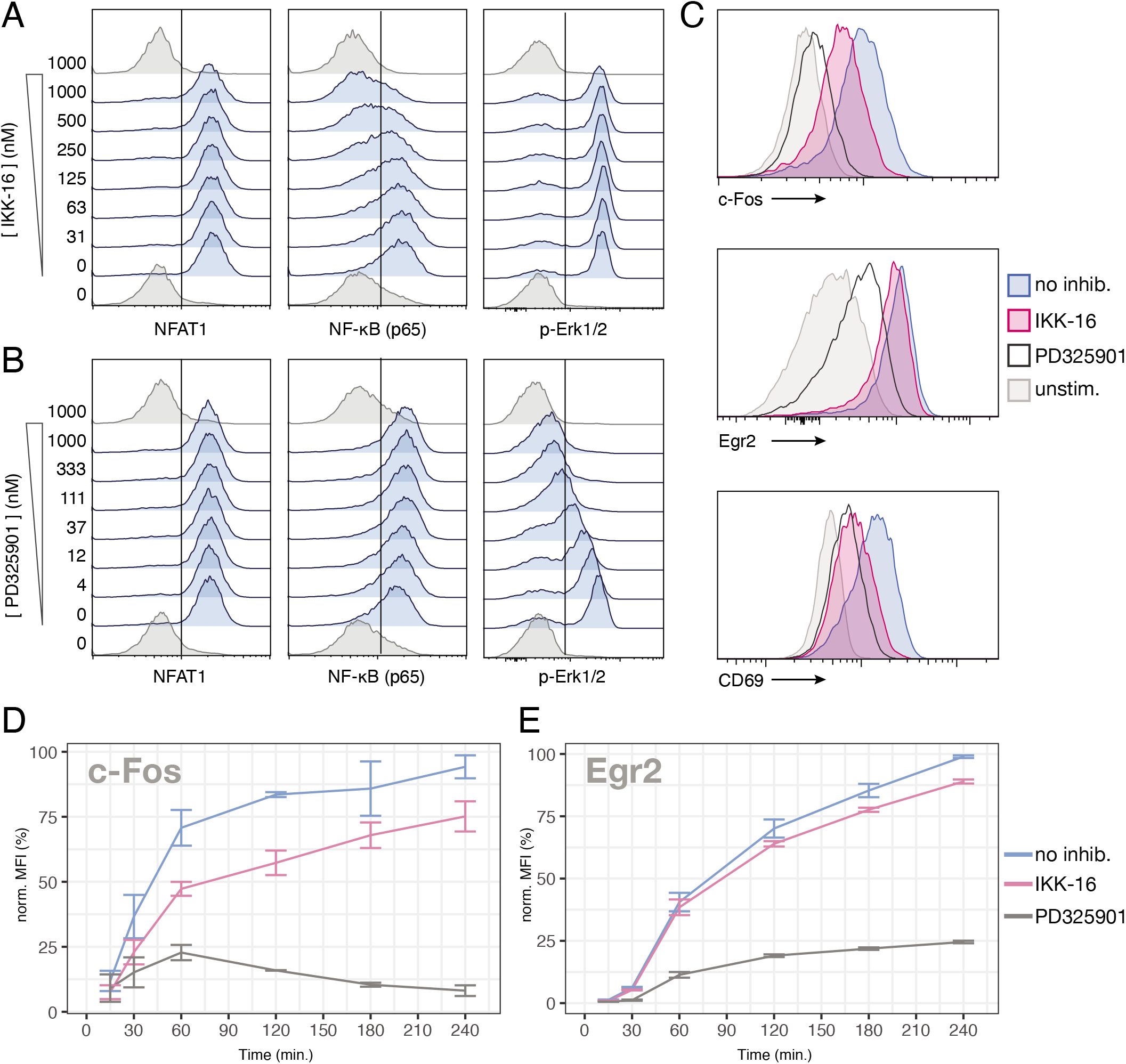
Specific NF-κB inhibition differentially reduces c-Fos protein accumulation. (**A-B**) Histograms depicting effect of titration of IKK-16 (A) or PD325901 (B) on NFAT1, NF-κB, and p-Erk1/2 activation after 1-hour simulation with N4 peptide. (**C**) Histograms demonstrating the effect of 500 nM IKK-16 or 1000 nM PD325901 on c-Fos, Egr2, and CD69 protein expression in OT-I cells stimulated with N4 peptide for 2 hours. (**D-E**) Line plots of depicting 500 nM IKK or 1000 nM PD325901 treatment on c-Fos (D) and Egr2 (E) protein accumulation over 4 hours stimulation with N4 peptide. Compiled are three separate experiments, error bars are s.e.m.

## DISCUSSION

Our results provide evidence for a hierarchy of signaling thresholds for pathways downstream of the TCR and ITK. NFAT, MAPK, and NF-κB signaling all exhibit varying patterns of activation in single naïve OT-I cells after stimulation with OVA peptide or altered OVA peptide ligands. We revealed NF-κB p65 translocation is uniquely sensitive to the quality of peptide:MHC interaction and ITK/RLK activity. Our findings also demonstrated that diminished NF-κB signaling due to ITK/RLK inhibition has transcriptional effects in stimulated T cells, in that NF-κB response genes and induced AP-1 gene family (*Fos, Fosl1, Fosb*) members are diminished in abundance compared to untreated cells.

Most models of TCR initiation describe receptor proximal signaling as digital and inherently noisy. In order for successful TCR signaling to result in changes to transcription factor activation in the nucleus, pathways must produce stable intermediates only after repeated and sustained engagements with peptide:MHC (5, 11, 61). TCR-induced MAPK activation has been shown to exhibit strong digital behavior due to positive feedback regulation of son of sevenless (SOS) which sustains active Ras (12, 61). To measure activation of MAPK signaling, we observed p-Erk1/2 fluorescence in single cells with flow cytometry. Consistent with previous reports, we measured strongly digital Erk activation in response to TCR ligation with peptide:MHC (11, 12). The fraction of cells positive for p-Erk after PRN694 treatment remained largely unchanged, but the p-Erk fraction was sensitive to peptide affinity (Fig. 1F). We expected p-Erk fluorescence to be sensitive to ITK activity because DAG production stabilizes SOS activation (12). However, it is possible that upstream phosphorylation of LAT after TCR triggering is the rate limiting step for MAPK activation rather than the ITK-dependent production of DAG. Further, initial experiments with *Itk-/-* mice reported decreased p-Erk in pooled whole CD8^+^ T cell lysates after stimulation with anti-CD3 antibody (21). In contrast, our experiments measure TCR responses after engagement with peptide:MHC ligands, which show markedly different kinetics compared to those elicited by α-CD3 stimulation.

A separate study previously reported NF-κB behaves digitally after TCR stimulation. Direct antibody-mediated stimulation of the TCR in C57BL/6 mouse CD4^+^ and CD8^+^ T cells or OVA peptide stimulation of CD4^+^ OT-II cells displayed evidence of digital NF-κB activation (13). In the current study, we provide a carefully tuned examination of physiological NF-κB responses in naïve CD8^+^ T cells. Given sufficiently strong peptide stimulation (native N4 OVA peptide) or PMA/Ionomycin, T cells can quickly and completely translocate p65 with similar kinetics as NFAT1, appearing “all or none” (Fig. 1C,E; 2A). Our results reveal however, the strength of TCR stimulation the amount of ITK activity produces intermediate, graded states of p65 activation during T cell priming.

As we observed that NF-κB signaling was specifically sensitive to PRN694 treatment, these findings highlighted a role for ITK in amplifying signals induced by weaker TCR inputs that could not sufficiently trigger NF-κB on their own, even under conditions that stimulate NFAT and Erk. Later during activation, after initial priming, combined TCR, CD28, and other co-stimulatory receptors become increasingly important in amplifying NF-κB signaling via PI3K (62). It may be advantageous for naive cells to limit NF-κB, by making it more difficult to trigger with TCR stimulus alone. Our results reveal however, weaker TCR stimulation or ITK/RLK inhibition with PRN694 produces intermediate states of p65 activation, which do not appear digital. Robust p65 nuclear translocation requires ITK and strong TCR interaction and NFAT1 and p-Erk remain digitally responsive during weaker TCR stimulation or PRN694 treatment. These results indicate discrete, analog levels of NF-κB activation during T cell priming.

NFAT translocation is exclusively dependent on SOCE activation of calcineurin, while NF-κB activation is layered with multiple signaling inputs (50, 63, 64). ITK and subsequent PLC-gamma-induced production of DAG activates NF-κB p65 through PKC-theta (17, 50). PLCgamma production of IP_3_ generates calcium flux and activates calmodulin and subsequently calcineurin, which dephosphorylates sequestered NFAT1. Calmodulin also activates calmodulin-dependent protein kinases (CAMK), which can stabilize the CARMA complex and ultimately assist in NF-κB activation (65). Thus, ITK influences activity of NF-κB by both DAG and IP_3_ production. Experiments with ionomycin alone stimulate modest NF-κB p65 activation in naïve T cells, albeit lessened compared to TCR stimulation with peptide:MHC (Fig. 1E; Fig. 2). We and others describe digital NFAT1 translocation and calcium flux in single cells, where a discrete threshold of activation governs an all-or-none response (Fig. 1C-D) (7, 9, 10). We attribute the sensitivity and analog qualities of NF-κB activation in response to ITK activity to the combinatorial effects of simultaneous Ca^2+^ and DAG signaling (Fig. 2). Indeed, higher NF-κB MFI correlates with “switched on” NFAT1 fluorescence, indicating that digital TCR initiation and Ca^2+^ flux may precede NF-κB activation (Fig. 1C, E, F).

One of the questions driving our transcriptomics and genomics experiments was to discern whether ITK activity, and broadly, strength of TCR signal, directs diverging transcriptional programs or rather tunes the abundance of transcripts within one activation-associated gene set. RNA-seq results revealed that ITK inhibition or weaker TCR interactions with lower affinity peptide:MHC induce similar genes as strong TCR signaling (Fig. 3). A recent single cell transcriptomic study, thoroughly concluded that peptide stimulated OT-I cells activate a single effector transcriptional program and resultant effector cells have similar cytolytic capacity, independent of TCR signal strength (46). With single cell RNA-seq analysis, the same study also showed weaker TCR signaling delays early gene transcription. We identified the temporally-induced early gene clusters within 2 hours of TCR stimulation, but did not detect similar shifts in transcriptional kinetics due to ITK inhibition or weaker peptide affinity, but this is likely a limitation of our pooled RNA-seq experiments. In contrast, our analyses of signaling events in naïve OT-I cells do show weaker stimulation and ITK inhibition delays onset of peak fractional NFAT responders and NF-κB is only appreciably detectable after 60 mins. Additionally, cytometric analysis of CD69, c-Myc, Egr2, and c-Fos display slowed expression when stimulated with weak peptide:MHC.

Cooperation of NFAT, AP-1, and NF-κB transcription factors is required for many aspects of optimal transcription of T cell activation programming, including production of cytokines like IL-2, which was one of the most differentially expressed genes due to TCR signal strength or PRN694 treatment after 2 hours (Supplemental Fig. 4). Early IL-2 signaling through the induced high-affinity IL-2 receptor CD25 helps maintain levels of c-Myc, while also lowering the apparent TCR signaling threshold (66, 67). Thus, CD8^+^ T cells that quickly transcribe large amounts of IL-2 can maximize their clonal expansion. The IL-2 promoter contains NF-κB binding sites and critical NFAT:AP-1 binding sites, where both partners are required for transcription (55). In ITK-inhibited cells or during weak signaling, we measure reduced NF-κB activation, diminished production of AP-1 transcripts (*Fos, Fosb, Fosl1*), and decreased Fos protein (Fig. 4). These conditions could contribute to slower production of IL-2 transcripts, which are among the most sensitive to the strength of TCR within 2 hours. There is evidence that T cells may continue to accumulate c-Fos protein after serial encounters with APCs during the early periods of T cell priming, effectively summing their cumulative duration of signaling (68, 69). TCR stimulation that induces efficient NFAT translocation, but not NF-κB, and only weakly induces AP-1 factors may suffer from low accumulation of early gene products and weak IL-2 production. Chromatin immunoprecipitation assays utilizing a constitutively active NFAT mutant, where NFAT is permanently nuclear, shows NFAT cannot bind the IL-2 promoter without its AP-1 binding partner; in the absence of AP-1, IL-2 transcription is abrogated (70).

A better understanding of the biology of T cell exhaustion is crucial in treating chronic illness and maximizing the efficacy of T cell immunotherapies, such as CAR-T. Recent work has identified NFAT as an important TCR-dependent regulator of T cell exhaustion phenotypes (70–72). NFAT can bind to the promoter of select exhaustion-associated genes without its partner AP-1 (70) and drives expression of NR4A family transcription factors which further maintain exhaustion programming (71, 72). Within the hierarchy of T cell signaling pathways we identify here, NFAT and MAPK appear to be more easily stimulated than NF-κB. Our data provide further evidence that CD8 T cells may signal through NFAT more easily, without complete downstream pathway activation, which could contribute to exhaustion phenotypes.

---

**Supplemental Figure 1.**
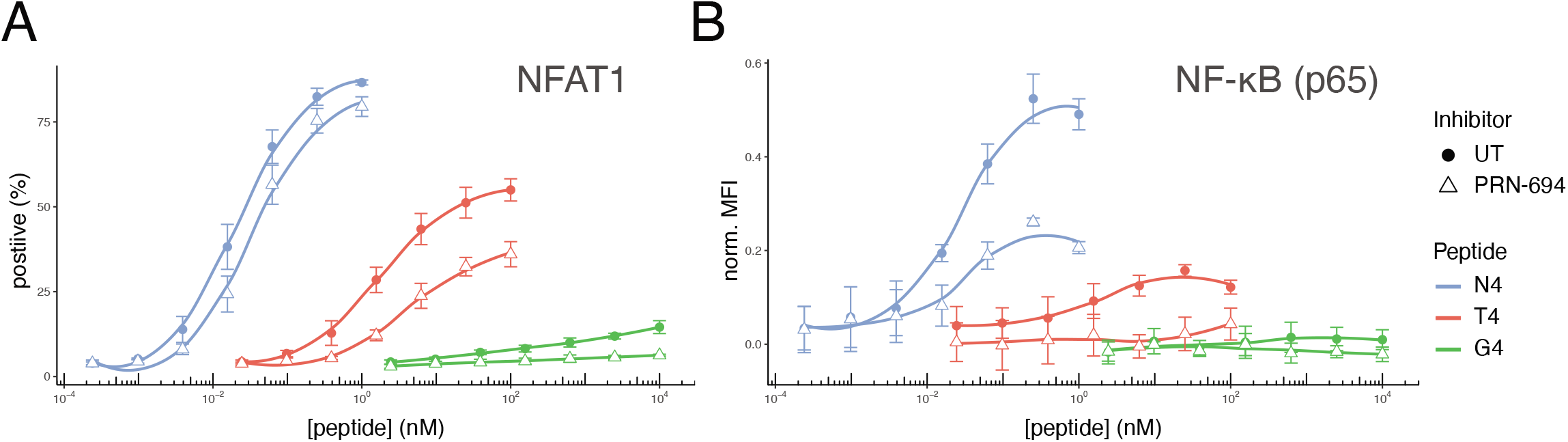
PRN694 differentially affects NF-κB and NFAT1 responses during variable peptide stimulation. (**A-B**) Line plots depicting NFAT1 or NF-κB (p65) activation in OT-I nuclei in response to 30 minute stimulation by WT splenocytes loaded with indicated doses of N4, T4, or G4 peptide, with or without treatment with 50 nM PRN694. Error bars indicate s.e.m.

**Supplemental Figure 2.**
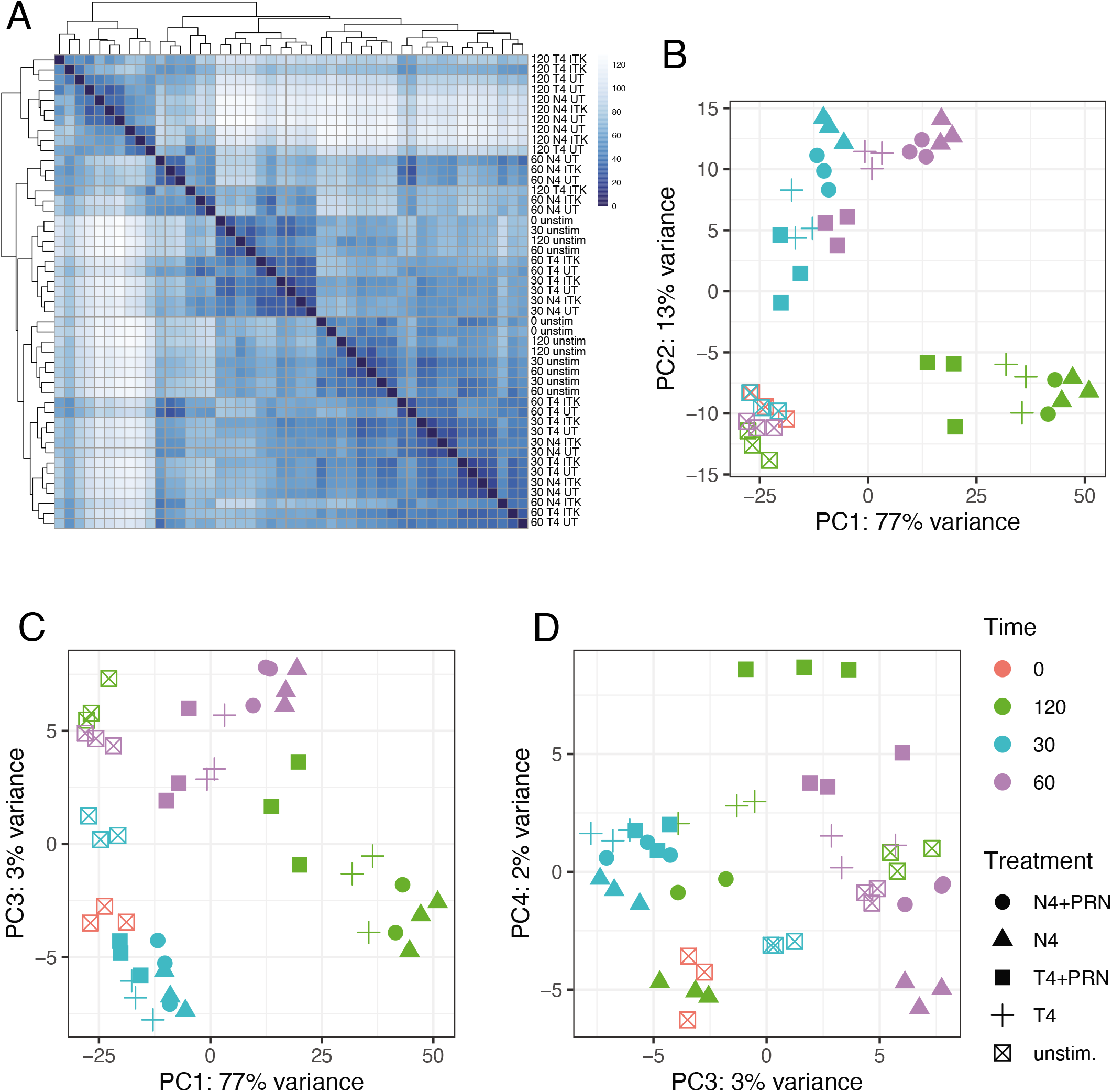
RNA-seq replicate correlation and principal component analysis. (**A**) Sample-to-sample correlation to compare RNA-seq replicates. (**B-D**) Principal component analysis of all RNA-seq replicates across all stimulation conditions.

**Supplemental Figure 3.**
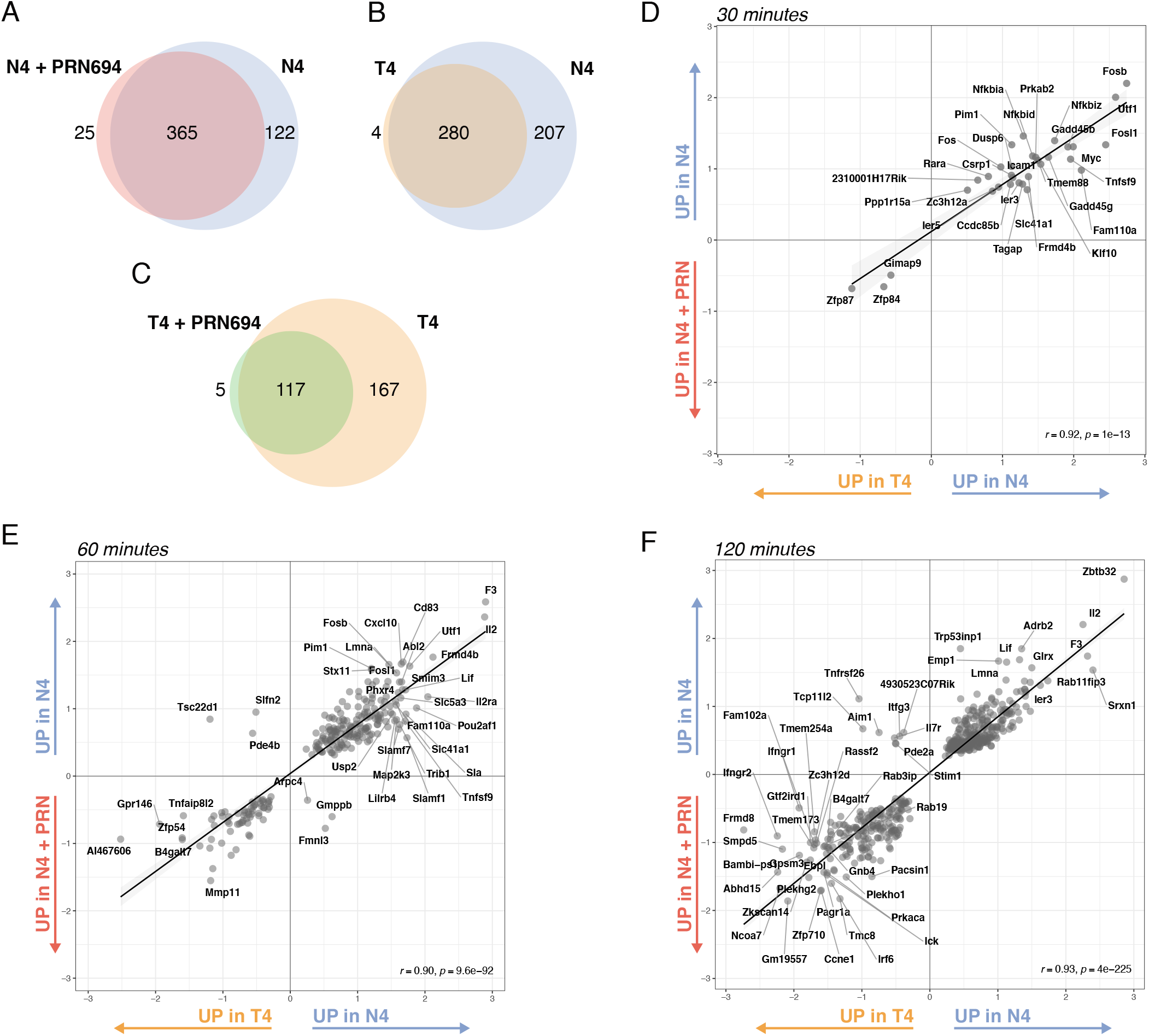
Graded TCR signaling induces a single transcriptional program. (**A-C**) Indicated overlap of induced genes identified in each tested TCR signaling condition. N4 = blue (*n*=487), T4 = orange (*n*=284), N4 + PRN694 = red (*n*=390), T4 + PRN694 = green (*n*=122). (**D-E**) The effect of stimulation with weaker affinity peptide vs treatment with PRN-964. Linear correlation of Log_2_ fold-change values determined by differential expression analysis of either N4/N4+PRN694 or N4/T4 comparisons at each time point.

**Supplemental Figure 4.**
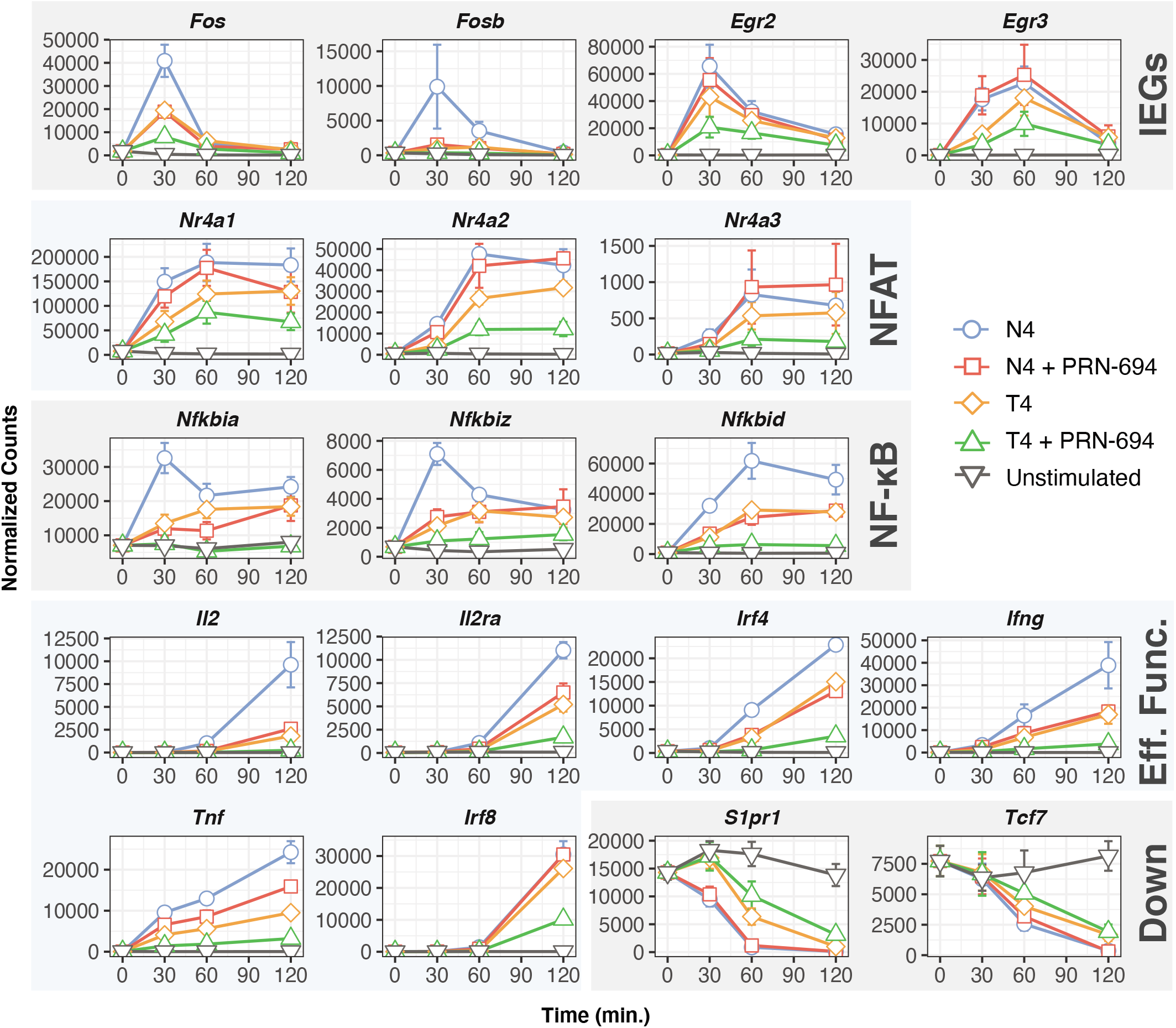
Example patterns of detected early transcripts. Line plots depicting normalized count data of selected gene transcripts over the course of the 2-hour RNA-seq experiment. Colors are consistent with other figures. Data represent three separate biological replicates. All genes presented are significantly differentially expressed compared to control at least at one time point. Error bars represent s.e.m.

**Supplemental Figure 5.**
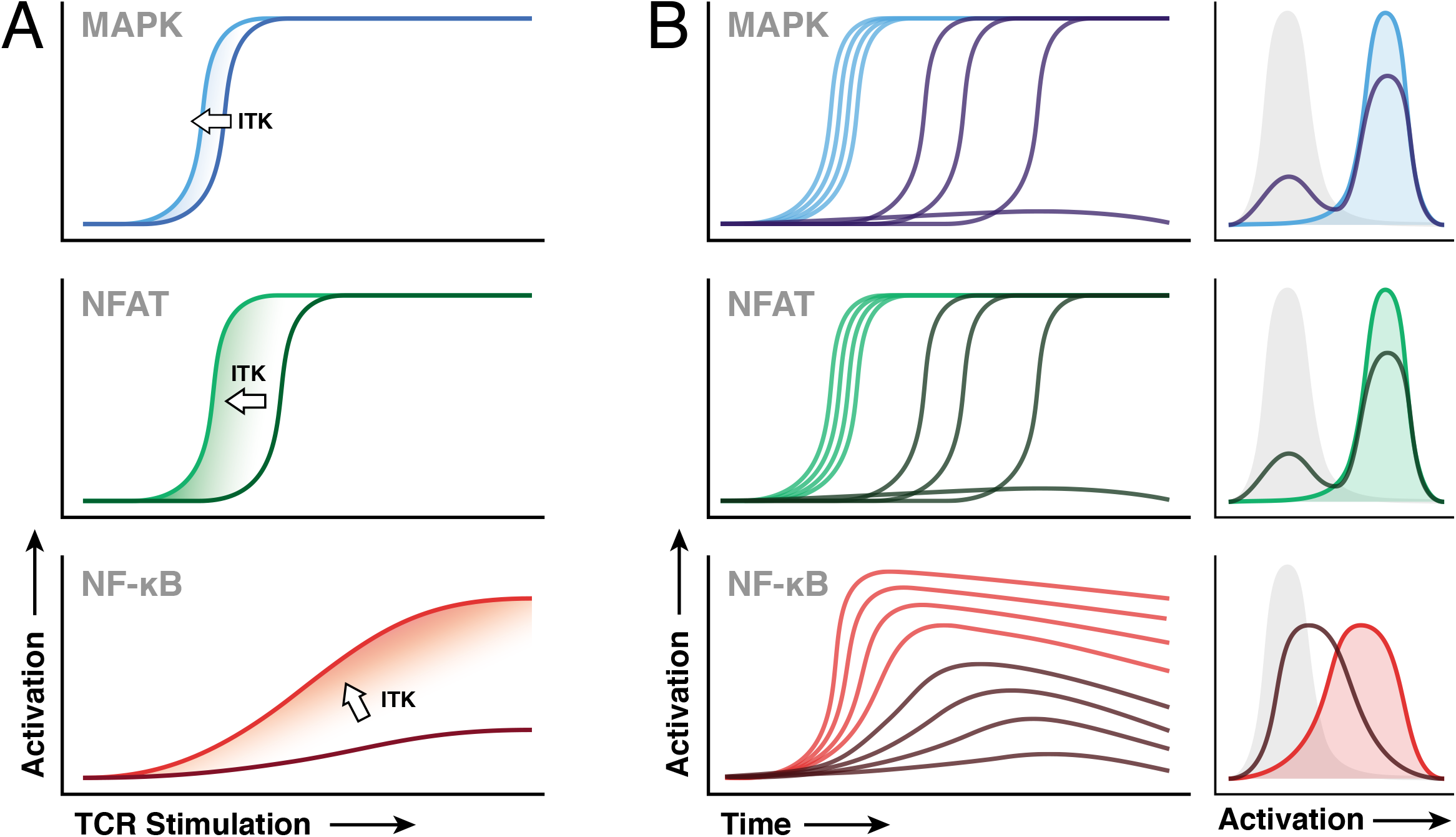
Data summary of TCR signal strength effects on activation of separate signal pathways. (**A**) Line plots depicting how ITK disproportionately shifts signaling thresholds. MAPK, NFAT, and NF-κB responses are drawn in blue, green, and, red respectively. Activation response when ITK is present is drawn in a lighter shade; activation without ITK is drawn in a darker shade. Direction of the shift due to presence of ITK is indicated with an arrow. “TCR Stimulation” input is plotted against the resultant degree of “Activation.” (**B**) Summary of TCR signal strength control of pathway activation within a stimulated population. Groups of curves representing single cell MAPK (blue), NFAT (green), and NF-κB (red) responses to either “strong” (light shade) or “weak” (dark shade) stimulation are presented over time. Histograms to the right represent average “snapshots” of pathway activation of the populations in the line plots.

## Notes

### Competing Interest Statement

The authors have declared no competing interest.

